# Visual Regularities Underlie Hierarchical Object Representations in the Human Brain and Self-supervised DNN

**DOI:** 10.1101/2025.08.31.672834

**Authors:** Qiande Zhao, Junhai Xu, Deying Li, Xia Wu, Kang Zhang, Congying Chu, Lingzhong Fan

## Abstract

NeuroAI develops the interplay of neuroscience and artificial intelligence, especially on visual processing. Human visual system organizes objects based on a representational hierarchy. However, it remains unclear whether this hierarchy arises from visual or semantic information. One hypothesis posits that the visual system is structured around statistical regularities of visual information. Here, we test this hypothesis using the THINGS datasets and pure-visual deep neural networks (DNN). We constructed a low-dimensional object space based on multiple abstract object properties, reflecting statistical patterns of visual regularities. By applying voxelwise encoding models, we identified clusters in the higher visual cortex based on their property tuning, and they were found to support specific object categories. These clusters serve as the middle level to reveal a property-cluster-object hierarchical organization. Subsequently, we investigated whether this hierarchical structure could be captured by a self-supervised DNN. Through activity similarity analysis, we mapped the brain clusters onto the DNN and independently found that the DNN’s clusters exhibited distinct property tuning and influenced the classification accuracy of corresponding object categories, mirroring the effects observed in the human brain. Our results demonstrate similar hierarchical structures in the human brain and self-supervised DNN, suggesting that the visual regularities shape neural architecture of visual system. This study highlights the great potential of neural computational model in neuroscience study.

**Index Terms** Visual Processing, Abstract Property, Hierarchical Representation, Self-supervised Visual DNN

## 1 INTRODUCTION

NeuroAI focuses on the interplay between neuroscience and artificial intelligence (AI). AI provides simulated platform to reveal computational mechanisms of brain and behavior, especially those which are hard to achieve with traditional neuroscience methods, while neuroscience gives insights into new path towards more powerful artificial general intelligence (AGI) [1, 2]. For decades, NeuroAI has achieved significant successes on visual processing. Humans recognize and interact with objects in the living environment relying on features such as color ^[3, 4]^, shape [5–7], animacy [7–10], and function [11–13]. Neuroscience has sought to unravel the neural mechanisms underlying how the brain organizes these features [14–19]. These neural mechanisms proceed the development of many powerful visual computational deep neural networks (DNNs), while visual DNNs in turn help to reveal computational principles of visual processing [20–22].

Humans naturally know that animals are animate while buildings are inanimate. This cognitive hierarchy is also observed in the visual system. Research in neuroscience has pinpointed a map of object category in the higher visual cortex (HVC), with discrete regions selectively responding to distinct object categories, such as face [23–25], body [26, 27], word [28], or scene [29–31]. Beyond this category selectivity, the arrangement of these regions is further aligned with many large-scaled representational maps of object dimensions (abstract concepts), including animacy [32, 33], real-world size [34, 35], aspect ratio [33, 36], or semantics [37–39]. These findings demonstrate a hierarchical organization that spans on different spatial scales. However, a critical question persists: does this cognitive hierarchy originate primarily from visual experience or from conceptual knowledge? Testing this question is complicated by the simultaneous and continuous interplay of bottom-up visual information and top-down semantic information in the human visual cortex, confounding efforts to isolate their individual contributions. A leading hypothesis in neuroscience proposes that the visual system is structured around naturally co-occurring patterns—statistical regularities of the visual environment—that predict behaviorally relevant outcomes [40–42]. To test this hypothesis, a computational model trained solely on visual inputs, e.g. self-supervised DNNs, paired with a comprehensive description of visual hierarchy, are essential to probe the origins of hierarchical organization of visual system.

Given these motivations, we leveraged the THINGS datasets—a rich repository of human neuroimaging and ratings of object properties across a large-scale image set—and the Topographic Deep Artificial Neural Network (TDANN), a self-supervised visual DNN trained with contrastive learning, to address these challenges [43–46]. In this study, we first constructed a low-dimensional object space using human ratings of 12 properties, which are critical abstract concepts to object representations (Fig. 1a). Using voxelwise encoding models, we determined representational maps of these properties in the human HVC, with distinct areas exhibiting specific tunings to these properties (Fig. 1b). By clustering these areas based on their property tunings, we obtained property-driven clusters that underpin the processing of preferred object categories. These clusters provide a middle level between property and category representational scales to form a comprehensive hierarchical organization. To test whether this property-cluster-object hierarchy emerge from only visual information, we projected these brain clusters onto the TDANN (Fig. 1c). Strikingly, the TDANN clusters displayed specific property tunings and influenced classification accuracy for corresponding object categories, paralleling effects observed in the human brain (Fig. 1d).

**Fig. 1.**
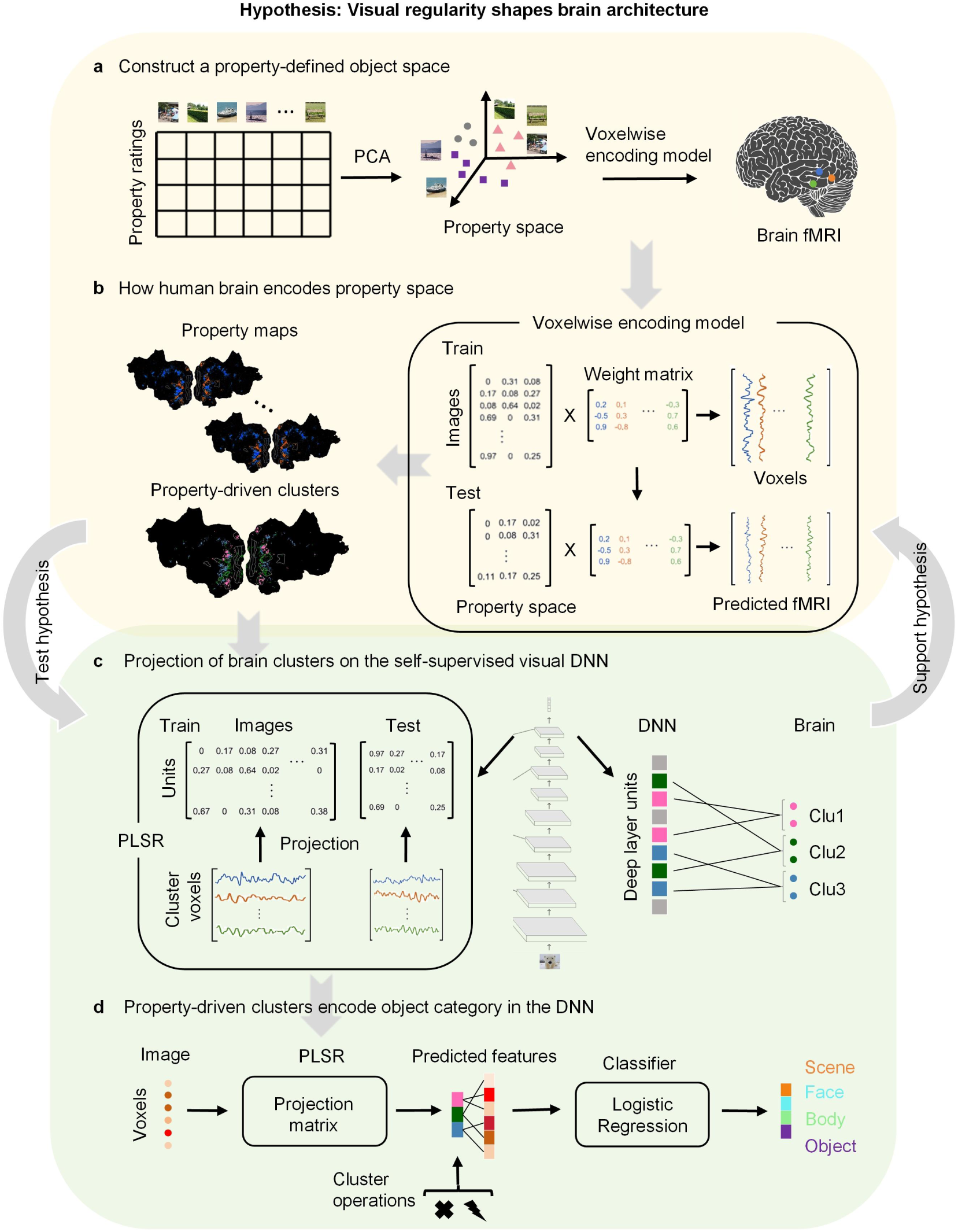
Framework for investigating property-driven hierarchical representations in the human brain and self-supervised DNN. **a**, Construction of property-defined object space. Principal component analysis (PCA) was performed on human cognitive ratings of abstract object properties to derive a low-dimensional object space. **b**, The voxelwise encoding model was then trained to analyze neural representations of this property space in human visual cortex. The weight matrix of the encoding model captures representational relationships between property space and neural responses. Property representation maps were visualized on cortical surfaces, and K-means clustering was applied to identify functionally distinct neural populations based on property tuning profiles. **c**, Property-driven brain clusters were projected onto a self-supervised visual DNN (TDANN) using partial least squares regression (PLSR). The projection matrix was used to classify TDANN units into cluster-selective (colored) and non-selective (gray) populations based on their correspondence with brain cluster responses. **d**, Functional characterization of the cluster in visual DNN. PLSR-predicted DNN features were systematically manipulated through either lesioning (X) or stimulating (lightning bolt) of cluster-selective units. The resulting feature perturbations were used to assess each cluster’s specific role in object representations through the downstream classification task.

Collectively, our main contributions are listed as follows: 1) We proposed property-driven clusters as middle level to unveil the hierarchy of object representations in HVC and self-supervised DNN model. 2) A computational model trained exclusively on visual information can naturally replicate the hierarchy of object representations observed in the human brain. 3) Through neuroAI framework, our findings bolster the hypothesis and highlight the power of visual regularities in shaping neural architecture. 4) Our study proceeds the understanding of visual processing and provides insights into more powerful visual AI model.

## 2 RELATED WORK

### 2.1 Representational hierarchy in the brain HVC

Visual system developed the hierarchical organization of object representations to support fast object recognition. Early research has identified several specialized regions in the higher visual cortex, such as the fusiform face area (FFA) [23–25] and the parahippocampal place area (PPA) [29–31], which are exceptionally responsive to specific object categories like faces or scenes separately. While, the discovery of such category-selective regions is limited, and representations of other object categories in the visual cortex remain largely uncharted [15, 47]. Moreover, these regions do not account for the entirety of the higher visual cortex, and functions of the remaining areas are not well understood [36, 48]. This has led to the study that the visual cortex is organized in a large scale along object dimensions reflecting abstract object properties, which were key to decide whether object was safe or not and how we could interact with it. These dimensional maps correspond with the arrangement of category regions [15, 32, 36]and that category regions exhibit specific profiles for many object dimensions [49–52]. These all demonstrate a hierarchical organization that spans on different spatial scales. While the gap between many dimensional maps and one-single category map still exists when we study this hierarchy. Therefore, we applied clustering algorithm to these dimensional maps to define dimension-tuning clusters to bridge the gap between above two level. Moreover, previous studies have been limited in scope, often focusing on one or two object concepts and employing contrast-based analyses on small image datasets [32, 33, 35]. Fortunately, THINGS datasets provide human ratings of many object properties across a large-scale image set and human neuroimaging data to these images. And voxel-wise encoding model can build linear mappings between multiple object properties and the fMRI signals of the whole brain voxels [37, 53–55]. These datasets and encoding methods would help to overcome above limitations.

### 2.2 Brain-aligned visual computational models

With the development of AI, the alignment of DNN and brain in functions and structures have been studied, especially on field of the visual cortex. Supervised DNNs, particularly those achieving human-level performance on visual classification tasks, can predict both behavioral and neural responses in the primate visual cortex [56–59]. Supervised DNNs encode some abstract object concepts, such as animacy, real-world size, that mirror those observed in the brain [50, 60]. Besides, these supervised DNNs also help to identify new dimensional maps in the monkey brain [36, 48], suggesting that object representations across brain scales form a unified hierarchical structure. However, when we study the origins of representational hierarchy of visual cortex, supervised DNNs are hard to tell the role of visual information and semantic information because of the prior labels of DNNs. While, recent work with self-supervised DNNs, trained exclusively on the visual data without labels, demonstrates that these models generate representational frameworks convergent with the visual stream at many levels [46, 52, 61, 62]. With pure visual information driven, self-supervised DNNs provide a new unifying platform for the computation of the human visual cortex. More importantly, these models provide the methods to unveil the origins of visual hierarchy which are hard to study in the human brain or supervised DNNs. Like the brain-score did [63–65], we mainly perform Partial Least Squared Regression (PLSR), a multivariate algorithm, to search for the counterpart of visual regions in the self-supervised DNNs based on their activity similarity.

Based on this, we later unveil the representational hierarchy of self-supervised DNNs and compare it with the brain.

## 3 Methods

### 3.1 THINGS datasets

The THINGS data is a multimodal neuroimage datasets that include fMRI and EEG data from 3 subjects viewing images, continuous human ratings of 12 object properties, and triplet odd-one-out behavior task data [43–45].

#### Neuroimaging data

The fMRI experiment scanned 3 subjects when viewing THINGS images. All magnetic resonance images (MRI) were collected using a 3 Tesla Siemens Magnetom Prisma scanner and a 32-channel head coil. fMRI was 2 mm isotropic resolution (60 axial slices, 2 mm slice thickness, no slice gap, matrix size 96×96, FOV = 192 × 192 mm, TR = 1.5 s, TE = 33ms, flip angle = 75°). Additional high-resolution T1-weighted and T2-weighted images were collected.

The MRI dataset was preprocessed with fMRIPrep [66]. The steps included slice time correction, head motion correction, susceptibility distortion correction, co-registration between functional and anatomical images, brain tissue segmentation and cortical surface reconstruction. The retinotopic mapping and functional localizer data were used to define retinotopic regions in the early visual cortex and masks of the category-selective regions in the higher visual cortex. Generalized linear models(GLM) were applied to estimate image-wise responses across sessions [67]. Finally, the responses matrix (image trial × whole brain voxels) is obtained for each subject.

#### Human property ratings

The THINGS datasets used crowdsourcing to generate a broad set of object property ratings, including manmadeness, preciousness, animacy, pleasantness, arousal, and real-world size, 12 properties in total [45]. For example, Animacy ratings for each object image were collected by presenting subjects with the respective noun and asking them to respond to the property ‘something that lives’ on a Likert scale.

Real-world size ratings for each object concept were obtained in the following steps. Subjects were asked to indicate the size of a target object on a continuous scale, defined by nine reference objects spanning the size range of all objects. Other concept properties, including grasp, hold, heavy, be moved, precious, pleasant, arousing, moves, natural, and manmade, follow similar rating procedures.

In our study, we employed mean human property ratings to construct an initial image property matrix encompassing 1854 image categories × 12 property dimensions. To align with our fMRI experimental design comprising 720 object categories, we subsequently derived a reduced property matrix (720 × 12) by indexing from the fMRI stimulus set. Recognizing substantial intercorrelations among certain property dimensions (e.g., heaviness and real-world size), we performed principal component analysis (PCA) on the property matrix. This dimensionality reduction yielded three principal components (PCs) that collectively accounted for more than 80% of the total variance. These PCs demonstrated strong correlations with the majority of original property dimensions and were consequently selected as our final image property features for subsequent analyses.

### 3.2 Voxelwise encoding model

We divided the whole fMRI data and related image property PCs into training and testing datasets in a 9:1 ratio randomly for each subject. The training set has 648 image categories, and the testing set has 72 image categories. We mapped the object property space to the cortex through the voxelwise linear encoding models based on the training set [37, 55]. For each voxel in the human brain, we modeled its response to an image as a linear combination of the property features in the object space.

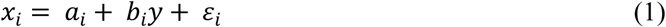

where *x_i_* is the beta value at the i-th voxel obtained from GLM analysis, *y* is the object property PCs in terms of a 3-dimensional column vector with each element corresponding to one axis (or feature) in the property space, *b_i_* is a row vector of regression coefficients, *a_i_* is the bias term, and *ε_i_* is the error or noise.

#### Training procedures

We have 648 object concepts beta values for each voxel of the human brain in the training set, which can be viewed as voxel responses across time t, with each time point relating to an imaging trial. It follows that the response of the i-th voxel at time t was expressed as Eq. (2)

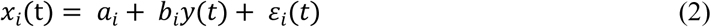

Least-squares estimation with L2-norm regularization was used to estimate the coefficients (*a_i_*; *b_i_*) given time samples of (*x_i_*; *y*). This is achieved by minimizing the loss function and each voxel is considered separately,

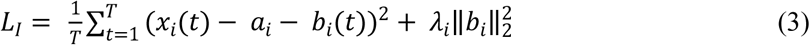

where T is the number of objects, and *λ_i_* is the regularization parameter for the i-th voxel. We used a 10-fold cross-validation to optimize the regularization parameter.

The statistical significance of each voxel was assessed based on a block-wise permutation test [68]. The training image set was divided into many blocks and the sequence of these blocks was randomly shuffled for 100,000 permutation trials. This would result in a null distribution. Then, we compared the original cross-validation without permutation to this null distribution. The one-sided p-value was calculated while testing the significance with FDR q<0.05. For more details about the statistical significance, please refer to [55].

#### Testing procedures

In this part, the generalization performance of the encoding model was assessed on the testing set. Image property PCs in the testing set were put into the trained encoding model, and the predicted fMRI responses to these 72 test images were generated, as indicated in the equation (4),

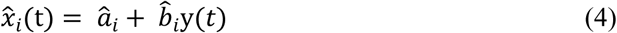

where y(t) is the vector of object property PCs across testing set images. Then, predicted voxel responses were correlated with actual responses to calculate the Pearson correlation as the model’s performance. The statistical significance procedure is similar to the operation in the training procedures and was applied to test the statistical significance of each voxel. The following analysis was based on the brain voxels, which can be significantly predicted by property space.

### 3.3 Visual hierarchy in the human brain

The Property representation maps were investigated through a systematic analysis of the encoding model weight matrix (voxels × property PCs). Cortical surface projections of individual PC weights were generated using pycortex [69], producing three distinct property maps. Additionally, an RGB-coded composite representation integrated all three PCs into a unified property space map for comprehensive visualization.

Voxels significantly predicted by the property PCs were subjected to K-means clustering, with optimal cluster number (k=3) determined through Calinski-Harabasz index evaluation.

The resulting clusters were visualized both in PC weight space and on cortical surface. Cluster-specific property tuning profiles were derived by averaging weights across constituent voxels for each cluster. Cluster-specific stimulus preferences were quantified by averaging voxel responses to object images within each cluster. Preferred stimuli were identified for each cluster as the top 10 images ranked by mean response magnitude, providing a quantitative measure of category selectivity.

### 3.4 Computational investigation of representational hierarchy in the pure-visual DNN model

The Topographic Deep Artificial Neural Network (TDANN) was developed to establish a unified framework for modeling functional organization across the early and higher-level visual stream [46]. Implemented using a ResNet-18 architecture, the model incorporates two fundamental optimization objectives during training. A contrastive loss function enforces metric learning in the latent space by minimizing representational distances between similar objects while maximizing separation for dissimilar objects. Concurrently, a spatial regularization term constrains inter-layer connectivity to maintain biological plausibility while reducing model complexity. This dual-optimization approach enables simultaneous prediction of object representations and simulation of topographic organization in both primary visual cortex (V1) and ventral temporal cortex (VTC), providing a powerful platform for investigating property-defined object space encoding mechanisms.

Image features were extracted from the VTC-like layer (layer 4.1) of a pre-trained TDANN model (model_final_checkpoint_phase199.torch) using stimuli from the THINGS datasets, with cortical mapping positions specified in the associated retinotopic initialization files. Both neural responses (images × voxels) and model activations (images × units) were randomly partitioned into train and test sets. Partial least squares regression (PLSR) was employed to predict TDANN features from corresponding neural responses in the train set, with model performance quantified by Pearson correlation between predicted and actual features in the test set. Comparative analyses were conducted using identical procedures for three additional supervised models (AlexNet FC layer, ResNet50 FC layer, CORNet-S IT layer) and an ablated TDANN variant trained without spatial constraints.

The PLSR projection matrix, capturing shared representational structure between biological and artificial systems (units × property-encoded voxels), was analyzed to establish unit-cluster correspondence. For each unit, mean projection weights were computed across voxels within each predefined brain cluster to obtain a triplet for each unit. Units were assigned to cluster-specific sets when meeting stringent criteria: maximal cluster weight > 0, and differential weighting (max - median) > (median - min). This classification yielded three cluster-selective unit populations and one non-selective set. Spatial distributions of these populations were visualized using the TDANN’s native cortical mapping framework, with unit selectivity profiles quantified through mean projection weights to each cluster.

To characterize property tuning profiles of each cluster in the VTC-like layer, we independently implemented a voxelwise encoding model that mapped property space to TDANN unit activations, mirroring our approach for biological neural data. The encoding weight matrix enabled quantification of property-specific responses for individual units. Cluster-level property tuning was derived by averaging weights across all units within each predefined cluster.

Given the observed anatomical overlap between brain clusters and known category-selective regions, we selected 36 exemplar images per category (scene, face, body, and manmade object) for training and 20 additional images per category for testing. A logistic regression classifier was trained on PLSR-predicted features to evaluate category classification. Two distinct manipulation protocols were implemented. First, lesion analyses were conducted through selective cluster manipulation. Cluster inclusion preserved activity in target clusters while nullifying all other units, whereas cluster exclusion suppressed target clusters while maintaining baseline activity elsewhere. Second, stimulation protocols systematically varied unit activations along a continuum from maximal inhibition (unit values minimized) to maximal excitation (unit values maximized), with non-target units held constant (zero).

## 4 RESULTS

### 4.1 Abstract property-defined object space

A low-dimensional object space was constructed utilizing multidimensional property ratings. Continuous-scale human cognitive ratings for 12 abstract object properties across 1,854 object concepts were obtained from the THINGS datasets [45], with dimensions including animacy, size, and other features previously established as critical for neural object representation. Distinct clustering patterns were observed along these dimensions: manufactured tools and biological organisms were found to occupy opposite extremes of the animacy axis, while systematic variation along the size axis was exhibited by objects ranging from small items (e.g., watches, spiders) to large entities (e.g., ships, whales).

Behavioral relevance among the twelve properties was revealed through initial analysis.

The predicted mutual exclusivity between ‘manmade’ and ‘natural’ dimensions was confirmed. These associations were systematically investigated through the computation of a representational dissimilarity matrix (RDM) across all property ratings. Three distinct clusters of highly correlated properties were identified (Fig. 2a). Dimensionality reduction was subsequently performed using principal component analysis (PCA), with three orthogonal components accounting for more than 80% of the variance being extracted (Fig. 2b).

**Fig. 2.**
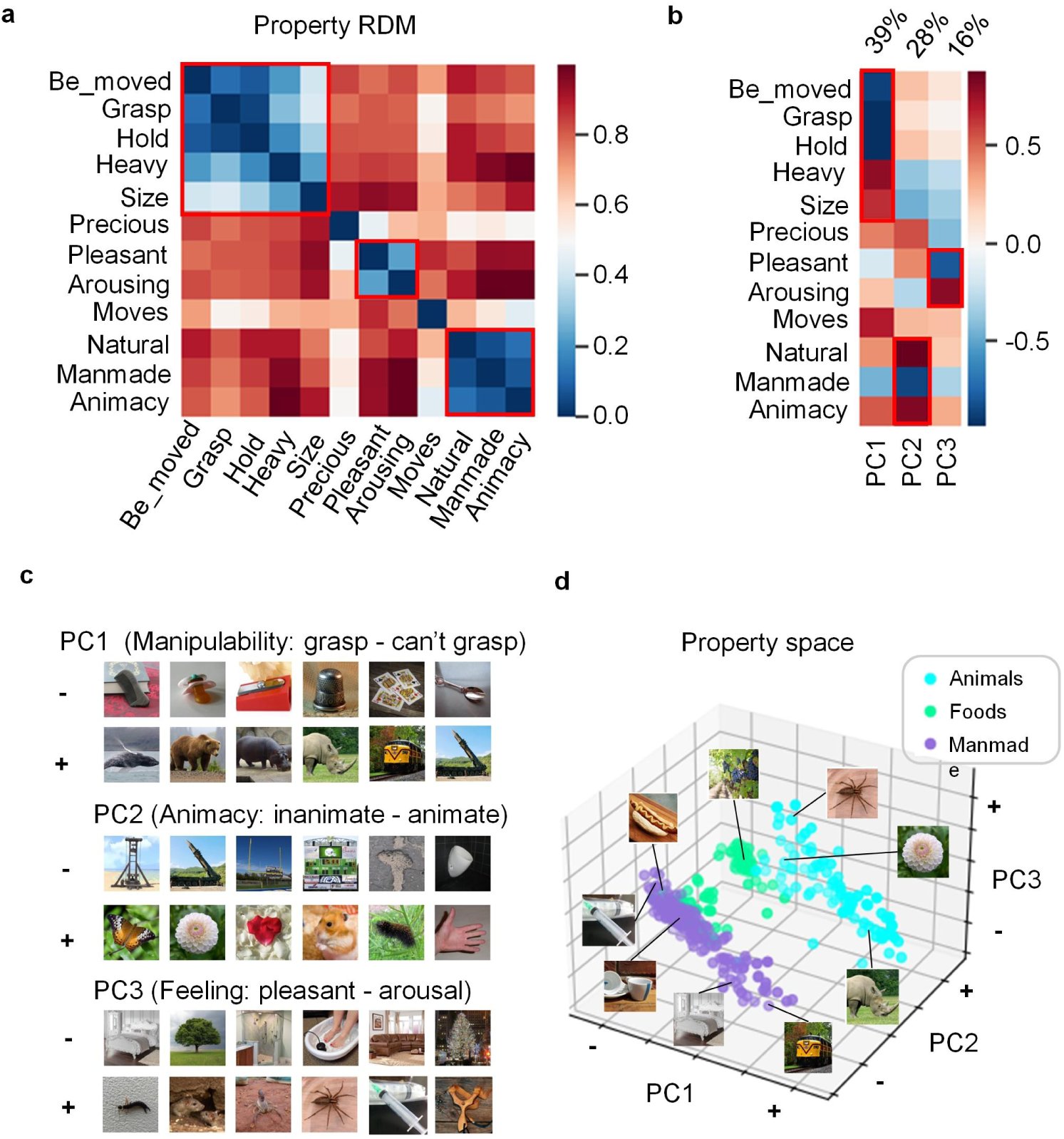
Construction and characterization of the property-defined object space. **a**, The representational dissimilarity matrix (RDM) was computed across 12 abstract property dimensions, with red rectangles highlighting groups of highly correlated properties. **b**, Principal component analysis (PCA) of object property ratings revealed that the first three principal components (PCs) collectively accounted for more than 80% of the variance. Correlation analysis demonstrated that each PC was predominantly associated with distinct property groups (highlighted in red). **c**, Representative image examples positioned at opposing ends (+/-) of each PC continuum. Based on their correlation patterns with original properties and characteristic image preferences, the three PCs were interpreted as representing manipulability, animacy, and feeling dimensions. **d**, Spatial distribution of object images within the three-dimensional PC space, color-coded by category membership (cyan: animals; light green: foods; purple: manmade objects). Selected exemplars are annotated to illustrate the organizational principles of this property-defined representational space.

Alignment between principal components (PCs) and property clusters was established through correlation analysis: PC1 was primarily associated with physical manipulability, PC2 with the animate-inanimate distinction, and PC3 with feeling. Validation of this dimensional structure was achieved through projection of category exemplars onto the PC-defined space (Fig. 2c-d). Clear differentiation along each dimension was observed: manipulable tools were localized to PC1- regions, while large, non-graspable entities were concentrated in PC1+ areas. Similarly, animate beings were predominantly located in PC2+ space, contrasting with inanimate objects in PC2- regions. Feeling differentiation was evidenced by the segregation of arousing objects (PC3+) from pleasing items (PC3-).

These findings demonstrate that a psychologically meaningful, low-dimensional object space can be defined by three principal dimensions: physical manipulability, animacy, and feeling. This organizational framework is shown to capture fundamental aspects of human object representation, with each dimension reflecting distinct but complementary aspects of cognitive processing.

### 4.2 Abstract property representation maps in HVC

The neural encoding of abstract property-defined object space was investigated through voxelwise linear encoding models trained to predict fMRI responses using object property PCs. Notably, for the representative subject (sub-01), the model predicted neural activity with statistical significance (q < 0.05, FDR corrected) predominantly in the higher visual cortex (HVC), including the medial/lateral occipital cortex and ventral temporal cortex (VTC) (Fig. 3a). This pattern was consistently replicated across another two subjects (Fig. S2a, S3a), confirming the essential role of HVC in abstract property representation.

**Fig. 3.**
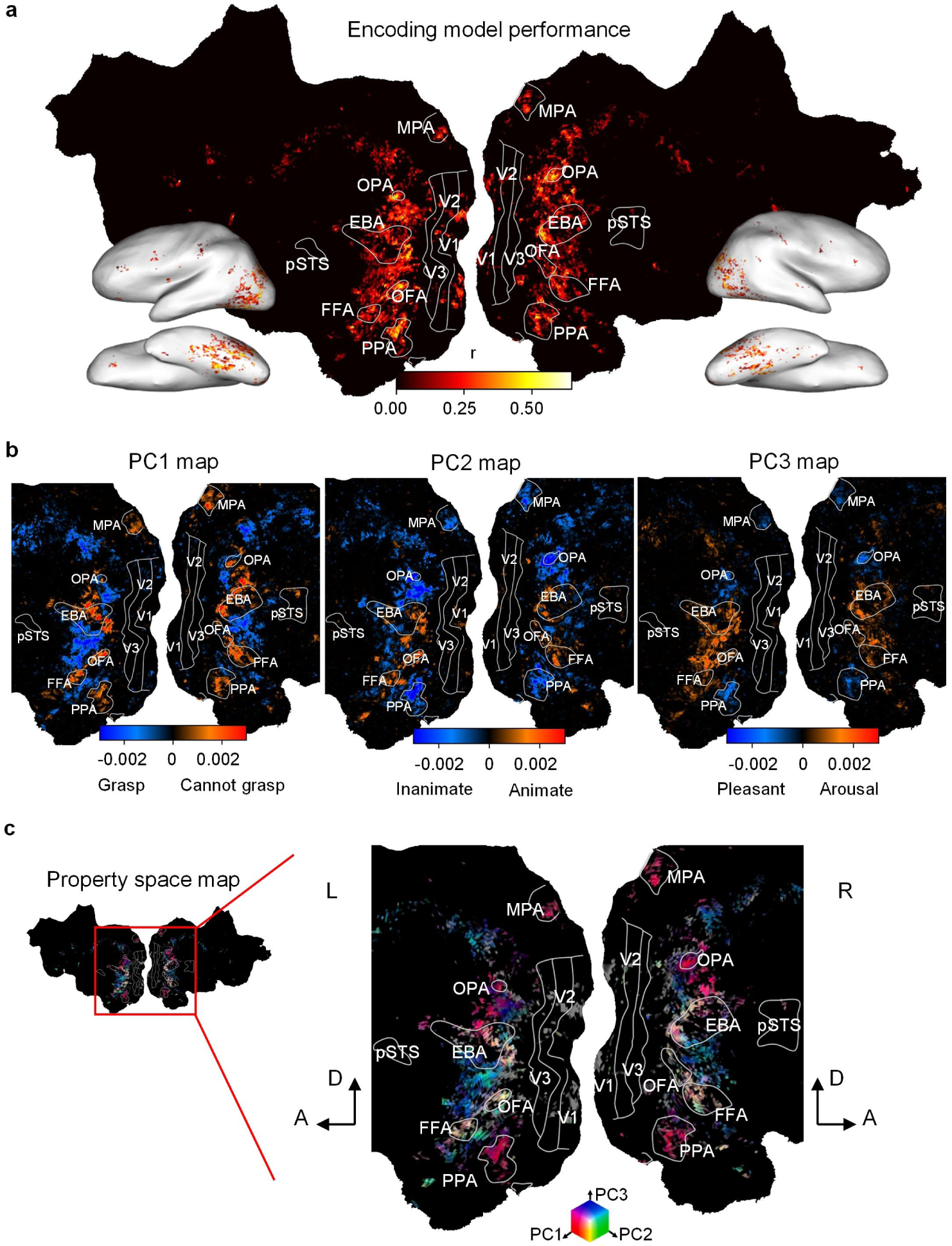
Property representation maps in human visual cortex (Sub-01). **a**, Prediction performance of voxelwise encoding models displayed on cortical surfaces. The map illustrates statistically significant Pearson correlation coefficients (q < 0.05, FDR corrected) between model-predicted and empirically measured neural responses to visual stimuli. **b**, Property representation maps generated by projecting PC-related weights from encoding models onto cortical surfaces. Color intensity reflects the magnitude of weight values for each property PC (manipulability, animacy, and affective dimensions). **c**, Integrated property map visualization using RGB code to represent the three property PCs. The inset provides a detailed view of the region indicated by the red rectangle, showing only voxels with statistical significance. All surface visualizations were generated using pycortex [69]. The early visual cortex includes V1, V2, and V3. Face-selective regions include FFA (Fusiform face area), OFA (Occipital face area), and pSTS (posterior superior temporal sulcus). Body-selective regions include EBA (Extrastriate body area). Scene-selective regions include PPA (parahippocampal place area), OPA (Occipital place area), and MPA (medial place area).

Following the establishment of HVC involvement, the cortical topography of property representations was examined through projection of encoding model weights onto the cortical surface (Fig. 3b). The resulting PC1 (manipulability) map revealed a striking dissociation: while ‘cannot grasp’ preferences (red) localized to scene, face, and body-selective regions, ‘grasp’ responses (blue) predominated in the lateral occipital cortex (LOC). This neural segregation is posited to reflect fundamental processing differences between manipulable artifacts and animate or scene stimuli. PC2 (animacy) patterns were precisely colocalized with face/body-selective areas, consistent with their established role in biological form processing. In contrast, PC3 (feeling dimension) responses exhibited inverse spatial distributions, with scene-selective regions showing a preference for pleasant stimuli (blue), while face, body, and object areas responded preferentially to arousing stimuli, suggesting distinct attentional modulation mechanisms for environmental versus interactive object processing. This topographic organization was robustly maintained across all examined subjects (Fig. S2b, S3b).

An integrated cortical map was generated through RGB codes of all three PC dimensions (Fig. 3c). Scene-selective regions were characterized by predominant PC1 weighting (red), while face and body processing areas (FFA/OFA/EBA) exhibited multiplexed property tuning. These findings collectively demonstrate that category-selective regions maintain distinct property preference profiles, revealing a systematic organizational principle governing object representation along manipulability, animacy, and feeling dimensions in human visual cortex.

### 4.3 Property-cluster-object hierarchical structures in HVC

The organization of property-category representations in the human visual cortex was investigated through K-means clustering of voxel-wise encoding weights. Quantitative evaluation of clustering metrics determined three optimal neural populations (k=3), whose distribution in PC-weight space demonstrated clear segregation (Fig. 4b). Distinct property tuning profiles were identified, with cluster 1 exhibiting preferential encoding of inanimate objects (PC2-), cluster 2 specializing in non-graspable and animate stimuli (PC1+), and cluster 3 selectively responding to graspable items (PC1-; Fig. 4c).

**Fig. 4.**
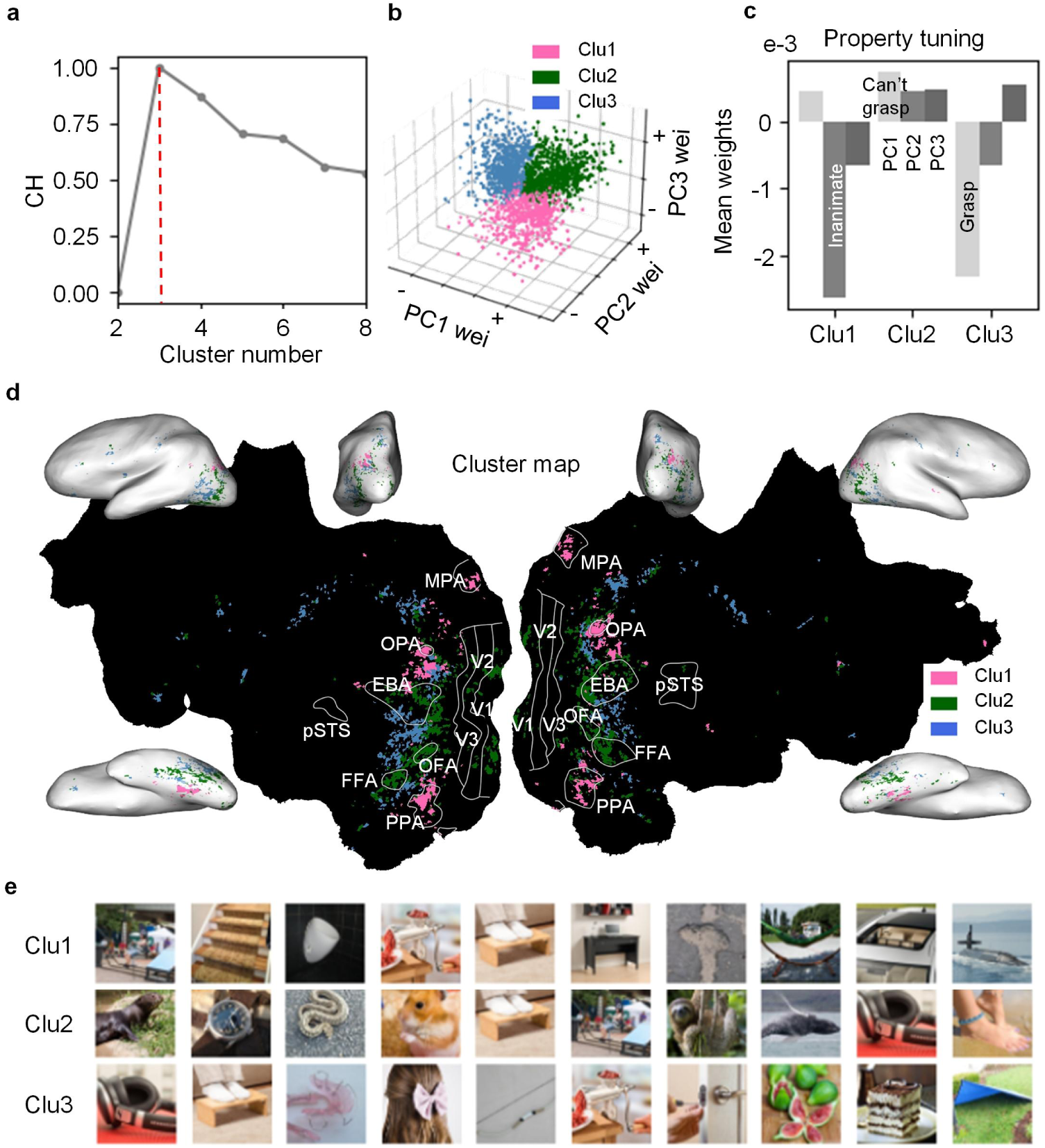
Property-driven cluster maps in the human visual cortex. **a**, Determination of optimal cluster number using the Calinski-Harabasz (CH) index, with maximum value indicating three distinct clusters. **b**, Spatial distribution of significantly predicted voxels in the property PC weight space, color-coded by cluster membership (hot pink, dark green, steel blue). **c**, Cluster-specific property tuning profiles derived by averaging PC-related encoding weights across constituent voxels for each cluster. **d**, Cortical map of property-driven clusters, maintaining consistent color scheme with panel B, demonstrating specialized topographic organization. **e**, Preferred object stimuli for each cluster, identified as the top ten images eliciting the strongest mean responses from cluster voxels.

The anatomical distribution of these clusters was found to precisely correspond with established functional regions (Fig. 4d). Specifically, cluster 1 (pink) was predominantly localized to scene-selective areas, including the parahippocampal place area, while cluster 2 (green) colocalized with face and body areas. In contrast, cluster 3 (blue) partially overlapped with object-selective regions (Fig. S1). This spatial organization was consistently replicated across additional subjects (Fig. S4) and quantitatively validated through cluster-category area ratio analysis (Fig. S5b). Further analysis of category preferences (Fig. 4e) revealed that cluster 1 responded maximally to scenes and big object ensembles, cluster 2 to animate stimuli, and cluster 3 to manipulable objects. These findings demonstrate that clusters serve as a milled level that systematically links specific property tuning profiles to anatomically distinct category regions and their preferred stimuli, and these all reveal a property-cluster-object hierarchical organization in the human HVC.

### 4.4 Clusters in the pure-visual model exhibited property tuning similar to the human brain

The TDANN model, trained exclusively on visual information, replicates key functional organization principles observed in primate V1 and ventral temporal cortex (VTC) [46]. Given this biological plausibility, we investigated how abstract object properties shape object representations in this nine-layer hierarchical model, with particular focus on its VTC-like layer (layer 4.1) due to its representational similarity to the human HVC (Fig. 5a).

**Fig. 5.**
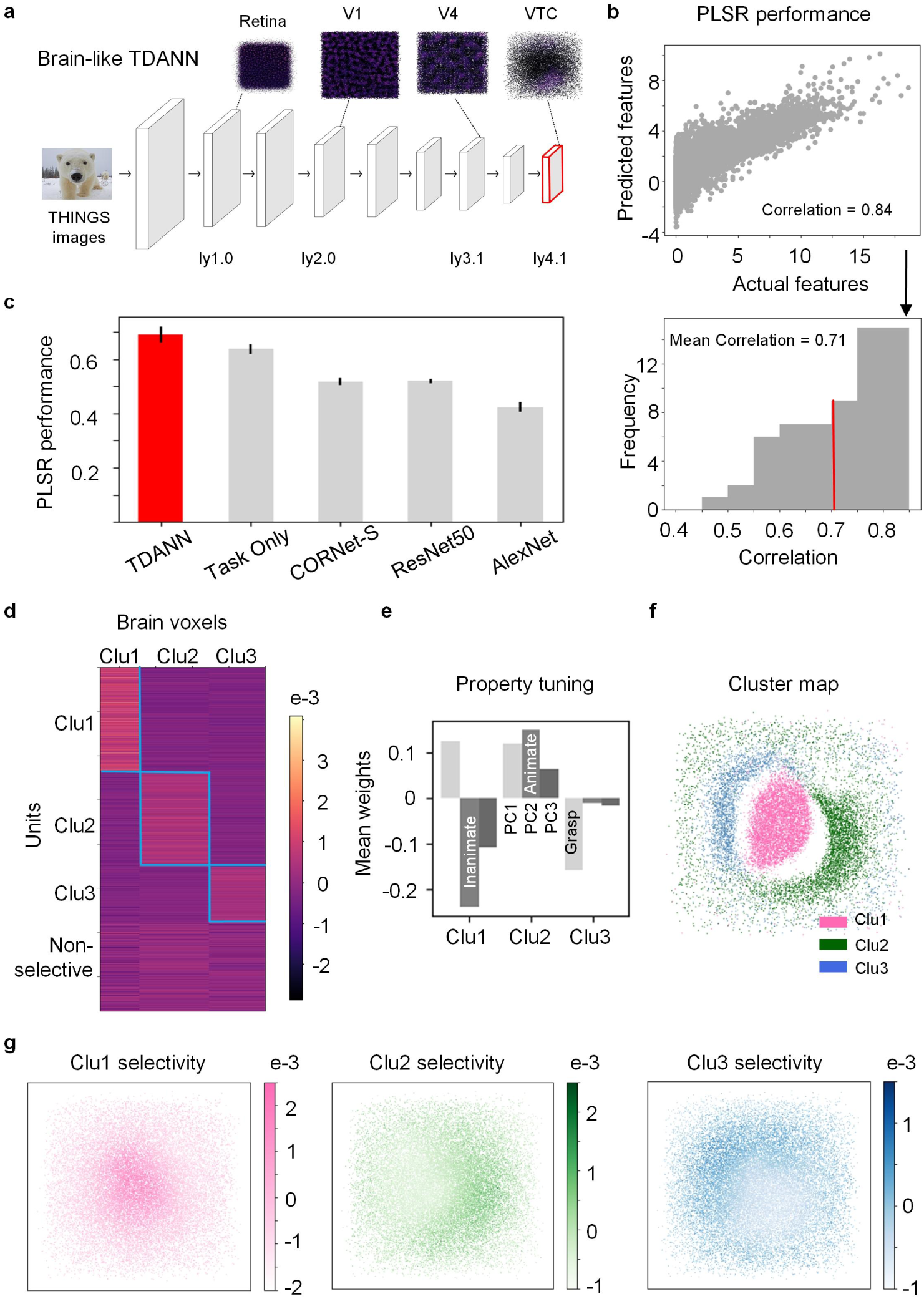
Projection of property-driven clusters onto the TDANN. **a**, Network architecture of the TDANN, featuring eight layers that simulate hierarchical visual processing from retina to ventral temporal cortex (VTC). Partial least squares regression (PLSR) was implemented to predict activations in the VTC-like layer (ly4.1) from cluster-specific neural responses in the training set. **b**, PLSR prediction performance for Sub-01. The top panel displays feature correlation for a representative test image, while the bottom panel shows the distribution of Pearson correlation coefficients across all test images (mean r = 0.71). **c**, Comparative PLSR performance across all subjects for the TDANN model versus control architectures, including a spatially unconstrained TDANN variant (Task Only) and three supervised visual models (CORNet-S, ResNet50, AlexNet). **d**, PLSR projection matrix revealing unit-cluster correspondence in the VTC-like layer. Units were classified as cluster-selective based on maximal mean projection weights to a given brain cluster exceeding statistical thresholds (see Methods). **e**, Property tuning profiles for cluster-specific units in the VTC-like layer, derived through independent voxelwise encoding models predicting unit responses from abstract property PCs. **f**, Spatial organization of cluster-selective units on the simulated cortical surface, maintaining color consistency with human cluster mappings. **g**, Quantification of unit preference for each brain cluster, expressed as mean projection weights from panel D. Higher values indicate stronger cluster preference, with color coding consistent across visualizations.

To quantitatively assess this similarity, we employed partial least squares regression (PLSR) to predict TDANN layer activations from human neural responses. Notably, the model achieved strong predictive performance (mean r = 0.71; Fig. 5b), which was consistently observed across all three subjects (Fig. 5c). Importantly, this performance significantly surpassed both spatially unconstrained self-supervised model (Task-Only) and three popular supervised visual DNNs, thereby establishing TDANN’s superior capacity for simulating cortical visual processing.

Building on these predictive relationships, we subsequently analyzed the PLSR projection matrix, which revealed distinct unit populations in the VTC-like layer corresponding to each brain cluster (Fig. 5d). Strikingly, these cluster-selective units exhibited property tuning profiles that closely mirrored their human neural counterparts (Fig. 5e), as further confirmed through independent voxelwise encoding models. Furthermore, spatial projection of brain clusters onto the simulated cortex (Fig. 5f) demonstrated segregated distribution patterns, wherein units within target clusters showed significantly higher selectivity magnitudes than non-target units (Fig. 5g). These results indicate that the TDANN model recapitulates property-specific representational geometry observed in human HVC, and does so through emergent abstract properties from the visual inputs.

### 4.5 Clusters in the pure-visual model affect the classification performance of objects similar to the human brain

To elucidate how object properties shape object representations, we first projected property-defined brain clusters onto the deep layers of the TDANN model. This approach therefore enabled systematic investigation of cluster-specific contributions to object classification through cluster-controlled computational manipulations (Fig. 1d). Specifically, we trained a four-category classifier (scene, face, body, and manmade object) using PLSR-predicted VTC-like layer features, subsequently applying two distinct experimental protocols to image features in the test set. Notably, cluster lesion experiments involved selective inclusion or exclusion of unit sets, while cluster stimulation systematically varied unit activations along an inhibition-excitation continuum (Fig. 6a). Non-selective units were excluded to ensure isolation of property-specific effects, with complete methodological details provided in the METHODS 3.4.

**Fig. 6.**
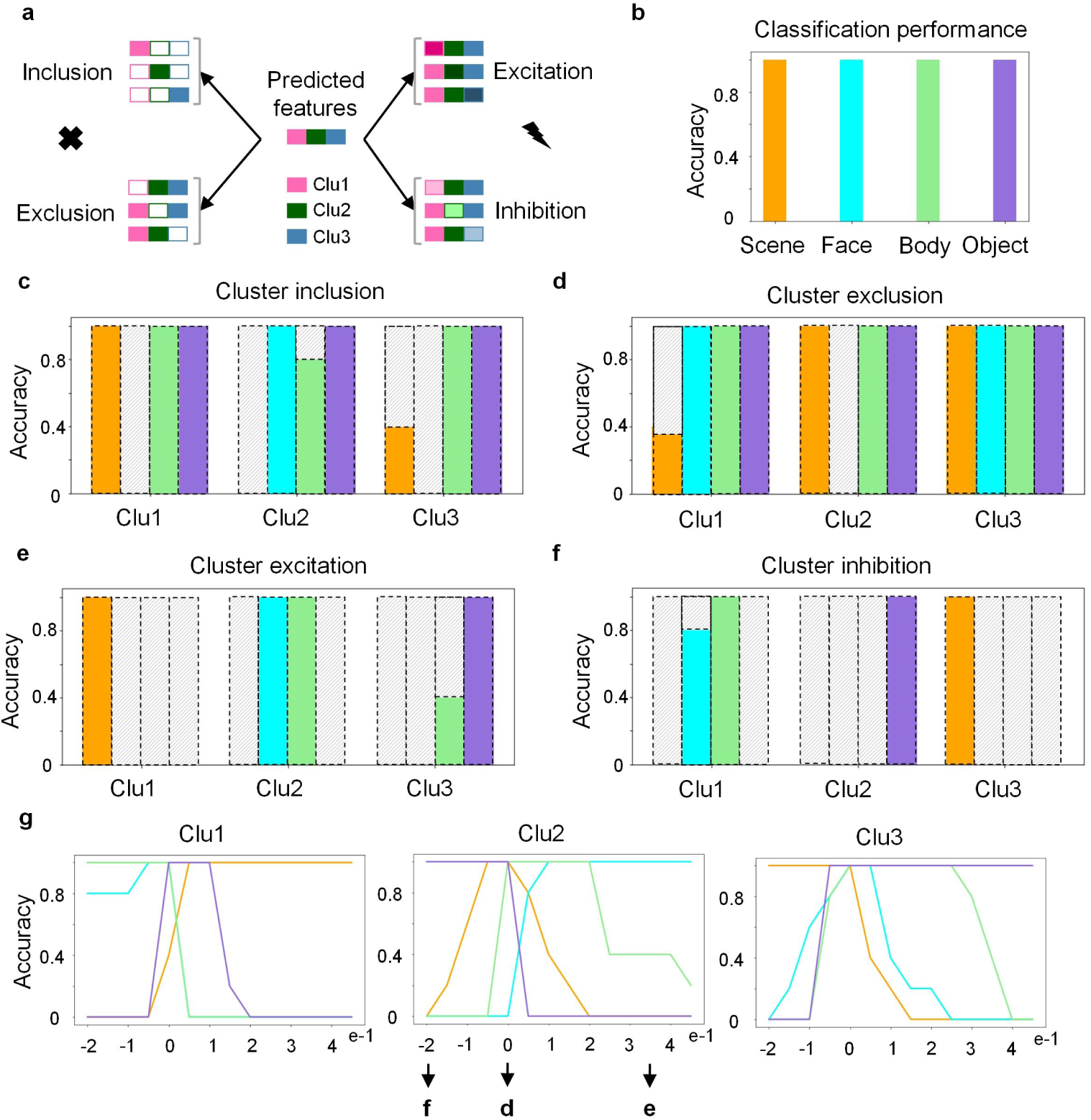
Analysis of object encoding by property-driven clusters in TDANN. **a**, Schematic of computational manipulations applied to PLSR-predicted features. Lesion operations (X) involved either inclusion (preserving target cluster units while zeroing others) or exclusion (zeroing target cluster units while preserving others). Stimulation (lightning bolt) systematically modulated unit activations along an excitation-inhibition continuum. Analyses focused exclusively on cluster-selective units (colored blocks), excluding non-selective components. Modified features were evaluated through a four-category classification task (scene: orange; face: cyan; body: light green; manmade object: purple). **b**, Baseline classification accuracy using unmodified PLSR-predicted features. **c**, Classification performance following cluster-specific inclusion manipulations. Dotted lines indicate baseline accuracy (b), colored regions show accuracy after manipulation, and gray hatched areas highlight accuracy differences. **d**, Classification accuracy following cluster-specific exclusion manipulations. **e**-**f**, Classification performance under maximal excitation (e) or inhibition (f) of cluster-specific units. **g**, Continuous accuracy changes during a stimulation gradient from full inhibition to full excitation, with marked points corresponding to inhibition, exclusion (zero), and excitation conditions shown in d-f.

The classifier initially achieved perfect baseline accuracy across all categories (Fig. 6b), establishing a robust foundation for subsequent manipulations. Lesion analyses revealed differential cluster specialization patterns. While all clusters contributed to manmade object and body representations, cluster 1 showed preferential involvement in scene processing and cluster 2 in face recognition (Fig. 6c). Similarly, exclusion manipulations demonstrated that cluster 1 lesions significantly impaired scene classification, whereas cluster 2 lesions completely abolished face recognition (Fig. 6d). The minimal impact of cluster 3 lesions strongly suggests functional redundancy in these object representations, indicating distributed processing mechanisms.

Building on these findings, stimulation experiments produced particularly striking category-specific biases. Maximal cluster 1 excitation universally biased classifications toward scenes, while cluster 2 excitation preferentially produced face and body categorizations, and cluster 3 excitation enhanced manmade object recognition (Fig. 6e). Remarkably, complementary inhibition protocols yielded precisely reciprocal effects. Cluster 1 inhibition enhanced face and body recognition, cluster 2 inhibition promoted manmade object classification, and cluster 3 inhibition biased responses toward scene categorization (Fig. 6f). Further, gradual stimulation generated continuous accuracy changes across the full spectrum of neural modulation, including inhibition, exclusion, and excitation conditions (Fig. 6g). These results demonstrate that property-defined clusters exhibit both specialized and distributed roles in object representation, closely recapitulating organizational principles observed in biological visual systems.

## 5 DISCUSSION

Our study first reveals a property-cluster-object hierarchical structure in the human higher visual cortex (HVC), constructed from a low-dimensional object space defined by abstract properties. By demonstrating that this hierarchical organization is mirrored in the Topographic Deep Artificial Neural Network (TDANN), a self-supervised model trained solely on visual information, we provide compelling evidence that the visual system is organized around statistical regularities in the visual world that predict behaviorally relevant outcomes. These findings align with the hypothesis introduced earlier, suggesting that naturally co-occurring patterns in the visual environment shape the neural architecture of object representation, independent of prior conceptual knowledge.

The critical role of abstract properties in object representation and recognition is well-established [49, 51, 70, 71]. Large-scale neural arrangements in the primate visual cortex are influenced by properties such as animacy, size, and aspect ratio, which reflect co-occurring patterns in the external world and their behavioral significance [51, 72–74]. Our study extends this understanding by integrating a more comprehensive set of 12 abstract properties, categorized into animacy-related, manipulability-related, and feeling-related dimensions (Fig. 2). Animacy-related properties distinguish natural from manmade objects, manipulability-related properties govern human-object interactions (e.g., tool use or food consumption) and correlate with object size, while affective properties arise from visual and behavioral experiences. Each property provides a principle for organizing objects in the brain, revealing the statistical object relationships embedded in the visual world. This multi-property approach overcomes the limitations of prior studies, which typically focused on one or two properties, offering a more holistic view of how the visual system structures object representations.

Our findings further demonstrate that traditional category-selective regions in the HVC exhibit preferences for specific profiles of behaviorally relevant properties, with their arrangements constrained by these abstract dimensions (Fig. 3). By parcellating the HVC into clusters based on property tunings, we identified distinct functional roles for each cluster, some overlapping with known category-selective regions and others encompassing areas with previously unclear functions (Fig. 4). These clusters can be viewed as a middle level to bridge the gap between continuous property dimensions and discrete category regions in the human HVC. This property-cluster-object hierarchy, illustrated in our results, suggests that abstract visual features extracted from environmental regularities and behavioral experience have the power to reveal hierarchical organization of visual systems. These features enable rapid object recognition and interaction, supporting efficient visual behavior. This organization aligns with and exceeds the hierarchical structure described in the Introduction, where representation maps of abstract concepts govern the arrangement of category-selective regions [49, 75–77].

A central challenge in visual neuroscience is disentangling the contributions of visual and semantic information to the hierarchical organization of the visual cortex, as these streams are intricately coupled in the human HVC [78]. To address this, we followed NueroAI framework to employ TDANN, a self-supervised model trained exclusively on the visual data, to test whether the property-cluster-object hierarchy emerges from visual information alone. Our results show a striking representational similarity between the human HVC and TDANN, with the model outperforming supervised deep neural networks (e.g., AlexNet, ResNet50, CORNet-S) in capturing brain-like property tunings (Fig. 5c). Using partial least squares regression (PLSR), we mapped brain clusters to TDANN and identified unit sets with selective responses to each cluster (Fig. 5d). Our independent encoding model shows that these TDANN clusters exhibited property tunings remarkably similar to those in the human brain, particularly for animacy- and manipulability-related dimensions (Fig. 5e). However, the model showed reduced selectivity for feeling-related properties, likely because these subjective emotional dimensions are not directly embedded in visual inputs, highlighting a limitation of purely visual models in capturing human-specific experiential factors.

To explore the functional roles of TDANN clusters, we conducted lesion and stimulation experiments on cluster-related image features and assessed their impact on object classification tasks. Each cluster exhibited preferences for specific object categories, interpretable through its property tunings, mirroring the functional specificity observed in the human HVC (Fig. 6). This convergence between the brain and TDANN underscores that a purely visual computational model can independently extract the property-cluster-object hierarchy, reinforcing the idea that visual regularities, rather than semantic knowledge, drive the organization of object representations. These findings build on the Introduction’s discussion of self-supervised DNNs converging with the hierarchical processing of the visual cortex, providing a direct test of the hypothesis that statistical regularities in the visual world shape neural architecture.

## 6 CONCLUSIONS

In conclusion, our study demonstrates that the property-cluster-object hierarchical structure of object representations in the human HVC can be naturally captured by a self-supervised visual model, supporting the hypothesis that the visual system is organized around co-occurring patterns in the visual environment that predict behaviorally relevant outcomes. By integrating a comprehensive property-defined object space with advanced computational modeling, we provide new insights into the origins of hierarchical object representation. These findings highlight the power of visual regularities in shaping neural and computational architectures, offering a foundation for developing more biologically inspired visual models and advancing our understanding of the visual system’s computational principles.

## DATA AND CODE AVAILABILITY

### Data

THINGS object concept and image database and metadata about human cognitive ratings of object properties are publicly available at https://osf.io/jum2f/, fMRI dataset is provided at https://doi.org/10.25452/Figshare.plus.c.6161151.v1. While TDANN model can be accessible at https://github.com/neuroailab/TDANN.

### Code

The code used for data processing, voxelwise encoding model, PLSR fitting, result analysis, and visualization in this study is publicly available as of the date of publication.

## ACKNOWLEDGE

This work was partially supported by STI2030-Major Projects (Grant No. 2021ZD0200200), the Natural Science Foundation of China (Grant Nos. 82472061, 82202253, 62250058), and Chongqing Science and Health Joint Medical Research Key Project (2025GGXM005). We also would like to thank Martin N. Hebart and Chris I Baker for sharing the THINGS and THINGSplus database and Eshed Margalit and Daniel L.K. Yamins for providing the TDANN model.

## DECALRE OF INTERESTS

Authors declare that they have no competing interests.

## REFERENCES

[1] A. Zador, S. Escola, B. Richards, B. Ölveczky, Y. Bengio, K. Boahen, M. Botvinick, D. Chklovskii, A. Churchland, C. Clopath, J. DiCarlo, S. Ganguli, J. Hawkins, K. Körding, A. Koulakov, Y. LeCun, T. Lillicrap, A. Marblestone, B. Olshausen, A. Pouget, C. Savin, T. Sejnowski, E. Simoncelli, S. Solla, D. Sussillo, A. S. Tolias, and D. Tsao, “ Catalyzing next-generation Artificial Intelligence through NeuroAI,” Nat Commun, vol. 14, no. 1, pp. 1597, Mar 22, 2023.

[2] T. Macpherson, A. Churchland, T. Sejnowski, J. DiCarlo, Y. Kamitani, H. Takahashi, and T. Hikida, “Natural and Artificial Intelligence: A brief introduction to the interplay between AI and neuroscience research,” Neural Netw, vol. 144, pp. 603–613, Dec, 2021.

[3] R. Shapley, and M. Hawken, “ Neural mechanisms for color perception in the primary visual cortex,” Curr Opin Neurobiol, vol. 12, no. 4, pp. 426–32, Aug, 2002.

[4] M. Li, N. Ju, R. Jiang, F. Liu, H. Jiang, S. Macknik, S. Martinez-Conde, and S. Tang, “Perceptual hue, lightness, and chroma are represented in a multidimensional functional anatomical map in macaque V1,” Prog Neurobiol, vol. 212, pp. 102251, May, 2022.

[5] A. A. Zeman, J. B. Ritchie, S. Bracci, and H. Op de Beeck, “Orthogonal Representations of Object Shape and Category in Deep Convolutional Neural Networks and Human Visual Cortex,” Sci Rep, vol. 10, no. 1, pp. 2453, Feb 12, 2020.

[6] V. Ayzenberg, and M. Behrmann, “ Does the brain’s ventral visual pathway compute object shape?,” Trends Cogn Sci, vol. 26, no. 12, pp. 1119–1132, Dec, 2022.

[7] F. Schmidt, M. Hegele, and R. W. Fleming, “Perceiving animacy from shape,” J Vis, vol. 17, no. 11, pp. 10, Sep 1, 2017.

[8] K. M. Jozwik, E. Najarro, J. J. F. van den Bosch, I. Charest, R. M. Cichy, and N. Kriegeskorte, “Disentangling five dimensions of animacy in human brain and behaviour,” Commun Biol, vol. 5, no. 1, pp. 1247, Nov 14, 2022.

[9] J. B. Ritchie, A. A. Zeman, J. Bosmans, S. Sun, K. Verhaegen, and H. P. Op de Beeck, “Untangling the Animacy Organization of Occipitotemporal Cortex,” J Neurosci, vol. 41, no. 33, pp. 7103–7119, Aug 18, 2021.

[10] S. Thorat, D. Proklova, and M. V. Peelen, “ The nature of the animacy organization in human ventral temporal cortex,” Elife, vol. 8, Sep 9, 2019.

[11] S. H. Johnson-Frey, “The neural bases of complex tool use in humans,” Trends Cogn Sci, vol. 8, no. 2, pp. 71–8, Feb, 2004.

[12] J. Almeida, A. Fracasso, S. Kristensen, D. Valerio, F. Bergstrom, R. Chakravarthi, Z. Tal, and J. Walbrin, “Neural and behavioral signatures of the multidimensionality of manipulable object processing,” Commun Biol, vol. 6, no. 1, pp. 940, Sep 14, 2023.

[13] E. L. Josephs, M. N. Hebart, and T. Konkle, “Dimensions underlying human understanding of the reachable world,” Cognition, vol. 234, pp. 105368, May, 2023.

[14] D. D. Cox, “ Do we understand high-level vision?,” Curr Opin Neurobiol, vol. 25, pp. 187–93, Apr, 2014.

[15] K. Grill-Spector, and K. S. Weiner, “ The functional architecture of the ventral temporal cortex and its role in categorization,” Nat Rev Neurosci, vol. 15, no. 8, pp. 536–48, Aug, 2014.

[16] J. J. DiCarlo, D. Zoccolan, and N. C. Rust, “ How does the brain solve visual object recognition?,” Neuron, vol. 73, no. 3, pp. 415–34, Feb 9, 2012.

[17] J. A. Collins, and I. R. Olson, “ Knowledge is power: How conceptual knowledge transforms visual cognition,” Psychonomic Bulletin & Review, vol. 21, no. 4, pp. 843–860, 2014.

[18] K. Grill-Spector, “The neural basis of object perception,” Curr Opin Neurobiol, vol. 13, no. 2, pp. 159–66, Apr, 2003.

[19] S. Bracci, and H. P. Op de Beeck, “ Understanding Human Object Vision: A Picture Is Worth a Thousand Representations,” Annu Rev Psychol, vol. 74, pp. 113–135, Jan 18, 2023.

[20] K. Kar, J. Kubilius, K. Schmidt, E. B. Issa, and J. J. DiCarlo, “Evidence that recurrent circuits are critical to the ventral stream’s execution of core object recognition behavior,” Nat Neurosci, vol. 22, no. 6, pp. 974–983, Jun, 2019.

[21] G. St-Yves, E. J. Allen, Y. Wu, K. Kay, and T. Naselaris, “ Brain-optimized deep neural network models of human visual areas learn non-hierarchical representations,” Nat Commun, vol. 14, no. 1, pp. 3329, Jun 7, 2023.

[22] D. L. Yamins, H. Hong, C. F. Cadieu, E. A. Solomon, D. Seibert, and J. J. DiCarlo, “ Performance-optimized hierarchical models predict neural responses in higher visual cortex,” Proc Natl Acad Sci U S A, vol. 111, no. 23, pp. 8619–24, Jun 10, 2014.

[23] N. Kanwisher, and G. Yovel, “The fusiform face area: a cortical region specialized for the ichBilSPlLconraTperception of faces,” os Rodi, vol. 361, no. 1476, pp. 2109–28, Dec 29, 2006.

[24] G. McCarthy, A. Puce, J. C. Gore, and T. Allison, “Face-specific processing in the human fusiform gyrus,” J Cogn Neurosci, vol. 9, no. 5, pp. 605–10, Fall, 1997.

[25] N. Kanwisher, J. McDermott, and M. M. Chun, “ The fusiform face area: a module in human extrastriate cortex specialized for face perception,” J Neurosci, vol. 17, no. 11, pp. 4302–11, Jun 1, 1997.

[26] M. V. Peelen, and P. E. Downing, “Selectivity for the human body in the fusiform gyrus,” J Neurophysiol, vol. 93, no. 1, pp. 603–8, Jan, 2005.

[27] P. E. Downing, Y. Jiang, M. Shuman, and N. Kanwisher, “A cortical area selective for visual processing of the human body,” Science, vol. 293, no. 5539, pp. 2470–3, Sep 28, 2001.

[28] A. Dȩbska, M. Wójcik, K. Chyl, G. Dziȩgiel-Fivet, and K. Jednoróg, “ Beyond the Visual Word Form Area - a cognitive characterization of the left ventral occipitotemporal cortex,” Front Hum Neurosci, vol. 17, pp. 1199366, 2023.

[29] R. Epstein, and N. Kanwisher, “A cortical representation of the local visual environment,” Nature, vol. 392, no. 6676, pp. 598–601, Apr 9, 1998.

[30] S. Nasr, N. Liu, K. J. Devaney, X. Yue, R. Rajimehr, L. G. Ungerleider, and R. B. Tootell, “Scene-selective cortical regions in human and nonhuman primates,” J Neurosci, vol. 31, no. 39, pp. 13771–85, Sep 28, 2011.

[31] G. K. Aguirre, E. Zarahn, and M. D’Esposito, “ An area within human ventral cortex sensitive to “building” stimuli: evidence and implications,” Neuron, vol. 21, no. 2, pp. 373–83, Aug, 1998.

[32] T. Konkle, and A. Caramazza, “Tripartite organization of the ventral stream by animacy and object size,” J Neurosci, vol. 33, no. 25, pp. 10235–42, Jun 19, 2013.

[33] D. D. Coggan, and F. Tong, “Spikiness and animacy as potential organizing principles of human ventral visual cortex,” Cereb Cortex, vol. 33, no. 13, pp. 8194–8217, Jun 20, 2023.

[34] S. Gabay, E. Kalanthroff, A. Henik, and N. Gronau, “ Conceptual size representation in ventral visual cortex,” Neuropsychologia, vol. 81, pp. 198–206, Jan 29, 2016.

[35] T. Konkle, and A. Oliva, “ A real-world size organization of object responses in occipitotemporal cortex,” Neuron, vol. 74, no. 6, pp. 1114–24, Jun 21, 2012.

[36] P. Bao, L. She, M. McGill, and D. Y. Tsao, “ A map of object space in primate inferotemporal cortex,” Nature, vol. 583, no. 7814, pp. 103–108, Jul, 2020.

[37] A. G. Huth, S. Nishimoto, A. T. Vu, and J. L. Gallant, “ A continuous semantic space describes the representation of thousands of object and action categories across the human brain,” Neuron, vol. 76, no. 6, pp. 1210–24, Dec 20, 2012.

[38] S. F. Popham, A. G. Huth, N. Y. Bilenko, F. Deniz, J. S. Gao, A. O. Nunez-Elizalde, and J. L. Gallant, “ Visual and linguistic semantic representations are aligned at the border of human visual cortex,” Nat Neurosci, vol. 24, no. 11, pp. 1628–1636, Nov, 2021.

[39] C. B. Martin, D. Douglas, R. N. Newsome, L. L. Man, and M. D. Barense, “Integrative and distinctive coding of visual and conceptual object features in the ventral visual stream,” Elife, vol. 7, Feb 2, 2018.

[40] H. S. Scholte, and E. H. F. de Haan, “ Beyond binding: from modular to natural vision,” TrendsCognSci, Apr 14, 2025.

[41] D. Kaiser, G. L. Quek, R. M. Cichy, and M. V. Peelen, “Object Vision in a Structured World,” Trends Cogn Sci, vol. 23, no. 8, pp. 672–685, Aug, 2019.

[42] A. Oliva, and A. Torralba, “The role of context in object recognition,” Trends Cogn Sci, vol. 11, no. 12, pp. 520–7, Dec, 2007.

[43] M. N. Hebart, A. H. Dickter, A. Kidder, W. Y. Kwok, A. Corriveau, C. Van Wicklin, and C. I. Baker, “THINGS: A database of 1,854 object concepts and more than 26,000 naturalistic object images,” PLoS One, vol. 14, no. 10, pp. e0223792, 2019.

[44] M. N. Hebart, O. Contier, L. Teichmann, A. H. Rockter, C. Y. Zheng, A. Kidder, A. Corriveau, M. Vaziri-Pashkam, and C. I. Baker, “THINGS-data, a multimodal collection of large-scale datasets for investigating object representations in human brain and behavior,” Elife, vol. 12, Feb 27, 2023.

[45] L. M. Stoinski, J. Perkuhn, and M. N. Hebart, “THINGSplus: New norms and metadata for the THINGS database of 1854 object concepts and 26,107 natural object images,” Behav Res Methods, vol. 56, no. 3, pp. 1583–1603, Mar, 2024.

[46] E. Margalit, H. Lee, D. Finzi, J. J. DiCarlo, K. Grill-Spector, and D. L. K. Yamins, “A unifying framework for functional organization in early and higher ventral visual cortex,” Neuron, vol. 112, no. 14, pp. 2435–2451.e7, Jul 17, 2024.

[47] R. Kiani, H. Esteky, K. Mirpour, and K. Tanaka, “ Object category structure in response patterns of neuronal population in monkey inferior temporal cortex,” J Neurophysiol, vol. 97, no. 6, pp. 4296–309, Jun, 2007.

[48] M. Yao, B. Wen, M. Yang, J. Guo, H. Jiang, C. Feng, Y. Cao, H. He, and L. Chang, “High-dimensional topographic organization of visual features in the primate temporal lobe,” Nat Commun, vol. 14, no. 1, pp. 5931, Sep 22, 2023.

[49] O. Contier, C. I. Baker, and M. N. Hebart, “ Distributed representations of behaviour-derived object dimensions in the human visual system,” Nat Hum Behav, Sep 9, 2024.

[50] F. R. Doshi, and T. Konkle, “Cortical topographic motifs emerge in a self-organized map of object space,” Sci Adv, vol. 9, no. 25, pp. eade8187, Jun 23, 2023.

[51] C. Magri, T. Konkle, and A. Caramazza, “The contribution of object size, manipulability, and stability on neural responses to inanimate objects, ” Neuroimage, vol. 237, pp. 118098, Aug 15, 2021.

[52] J. S. Prince, G. A. Alvarez, and T. Konkle, “Contrastive learning explains the emergence and function of visual category-selective regions,” Sci Adv, vol. 10, no. 39, pp. eadl1776, Sep 27, 2024.

[53] A. LeBel, S. Jain, and A. G. Huth, “ Voxelwise Encoding Models Show That Cerebellar Language Representations Are Highly Conceptual,” J Neurosci, vol. 41, no. 50, pp. 10341–10355, Dec 15, 2021.

[54] A. Mahmoudi, S. Takerkart, F. Regragui, D. Boussaoud, and A. Brovelli, “ Multivoxel pattern analysis for FMRI data: a review,” Comput Math Methods Med, vol. 2012, pp. 961257, 2012.

[55] Y. Zhang, K. Han, R. Worth, and Z. Liu, “Connecting concepts in the brain by mapping cortical representations of semantic relations,” Nat Commun, vol. 11, no. 1, pp. 1877, Apr 20, 2020.

[56] C. F. Cadieu, H. Hong, D. L. Yamins, N. Pinto, D. Ardila, E. A. Solomon, N. J. Majaj, and J. J. DiCarlo, “ Deep neural networks rival the representation of primate IT cortex for core visual object recognition,” PLoS Comput Biol, vol. 10, no. 12, pp. e1003963, Dec, 2014.

[57] R. M. Cichy, A. Khosla, D. Pantazis, A. Torralba, and A. Oliva, “Comparison of deep neural networks to spatio-temporal cortical dynamics of human visual object recognition reveals hierarchical correspondence,” Sci Rep, vol. 6, pp. 27755, Jun 10, 2016.

[58] K. R. Storrs, T. C. Kietzmann, A. Walther, J. Mehrer, and N. Kriegeskorte, “Diverse Deep Neural Networks All Predict Human Inferior Temporal Cortex Well, After Training and Fitting,” J Cogn Neurosci, vol. 33, no. 10, pp. 2044–2064, Sep 1, 2021.

[59] T. Horikawa, and Y. Kamitani, “ Generic decoding of seen and imagined objects using hierarchical visual features,” Nat Commun, vol. 8, pp. 15037, May 22, 2017.

[60] K. Dobs, J. Martinez, A. J. E. Kell, and N. Kanwisher, “Brain-like functional specialization emerges spontaneously in deep neural networks,” Sci Adv, vol. 8, no. 11, pp. eabl8913, Mar 18, 2022.

[61] I. Higgins, L. Chang, V. Langston, D. Hassabis, C. Summerfield, D. Tsao, and M. Botvinick, “ Unsupervised deep learning identifies semantic disentanglement in single inferotemporal face patch neurons,” Nat Commun, vol. 12, no. 1, pp. 6456, Nov 9, 2021.

[62] T. Konkle, and G. A. Alvarez, “A self-supervised domain-general learning framework for human ventral stream representation, ” Nat Commun, vol. 13, no. 1, pp. 491, Jan 25, 2022.

[63] M. Schrimpf, J. Kubilius, H. Hong, N. J. Majaj, R. Rajalingham, E. B. Issa, K. Kar, P. Bashivan, J. Prescott-Roy, and F. J. B. Geiger, “ Brain-score: Which artificial neural network for object recognition is most brain-like?,” pp. 407007, 2018.

[64] S. Nonaka, K. Majima, S. C. Aoki, and Y. Kamitani, “ Brain hierarchy score: Which deep neural networks are hierarchically brain-like?,” iScience, vol. 24, no. 9, 2021.

[65] M. Schrimpf, J. Kubilius, M. J. Lee, N. A. Ratan Murty, R. Ajemian, and J. J. DiCarlo, “ Integrative Benchmarking to Advance Neurally Mechanistic Models of Human Intelligence,” Neuron, vol. 108, no. 3, pp. 413–423, Nov 11, 2020.

[66] O. Esteban, C. J. Markiewicz, R. W. Blair, C. A. Moodie, A. I. Isik, A. Erramuzpe, J. D. Kent, M. Goncalves, E. DuPre, M. Snyder, H. Oya, S. S. Ghosh, J. Wright, J. Durnez, R. A. Poldrack, and K. J. Gorgolewski, “fMRIPrep: a robust preprocessing pipeline for functional MRI,” Nat Methods, vol. 16, no. 1, pp. 111–116, Jan, 2019.

[67] K. J. Friston, A. P. Holmes, J. B. Poline, P. J. Grasby, S. C. Williams, R. S. Frackowiak, and R. Turner, “Analysis of fMRI time-series revisited,” Neuroimage, vol. 2, no. 1, pp. 45–53, Mar, 1995.

[68] D. Adolf, S. Weston, S. Baecke, M. Luchtmann, J. Bernarding, and S. Kropf, “Increasing the reliability of data analysis of functional magnetic resonance imaging by applying a new blockwise permutation method,” Front Neuroinform, vol. 8, pp. 72, 2014.

[69] J. S. Gao, A. G. Huth, M. D. Lescroart, and J. L. Gallant, “Pycortex: an interactive surface visualizer for fMRI,” Front Neuroinform, vol. 9, pp. 23, 2015.

[70] M. N. Hebart, C. Y. Zheng, F. Pereira, and C. I. Baker, “ Revealing the multidimensional mental representations of natural objects underlying human similarity judgements,” Nat Hum Behav, vol. 4, no. 11, pp. 1173–1185, Nov, 2020.

[71] P. J. Lang, M. M. Bradley, J. R. Fitzsimmons, B. N. Cuthbert, J. D. Scott, B. Moulder, and V. Nangia, “ Emotional arousal and activation of the visual cortex: an fMRI analysis,” Psychophysiology, vol. 35, no. 2, pp. 199–210, Mar, 1998.

[72] D. Pitcher, L. Charles, J. T. Devlin, V. Walsh, and B. Duchaine, “Triple dissociation of faces, bodies, and objects in extrastriate cortex,” Curr Biol, vol. 19, no. 4, pp. 319–24, Feb 24, 2009.

[73] S. Bracci, and M. V. Peelen, “ Body and object effectors: the organization of object representations in high-level visual cortex reflects body-object interactions,” J Neurosci, vol. 33, no. 46, pp. 18247–58, Nov 13, 2013.

[74] B. Sorscher, S. Ganguli, and H. Sompolinsky, “Neural representational geometry underlies few-shot concept learning,” Proc Natl Acad Sci USA, vol. 119, no. 43, pp. e2200800119, Oct 25, 2022.

[75] M. A. Cohen, G. A. Alvarez, K. Nakayama, and T. Konkle, “ Visual search for object categories is predicted by the representational architecture of high-level visual cortex,” J Neurophysiol, vol. 117, no. 1, pp. 388–402, Jan 1, 2017.

[76] Groen, II, E. H. Silson, and C. I. Baker, “Contributions of low- and high-level properties to neural processing of visual scenes in the human brain,” Philos Trans R Soc Lond B Biol Sci, vol. 372, no. 1714, Feb 19, 2017.

[77] D. D. Coggan, D. H. Baker, and T. J. Andrews, “Selectivity for mid-level properties of faces and places in the fusiform face area and parahippocampal place area,” Eur J Neurosci, vol. 49, no. 12, pp. 1587–1596, Jun, 2019.

[78] A. Y. Wang, K. Kay, T. Naselaris, M. J. Tarr, and L. Wehbe, “ Better models of human high-level visual cortex emerge from natural language supervision with a large and diverse dataset,” Nau Machine Intelligence, vol. 5, no. 12, pp. 1415–1426, 2023.

